# SARS-CoV-2 exhibits intra-host genomic plasticity and low-frequency polymorphic quasispecies

**DOI:** 10.1101/2020.03.27.009480

**Authors:** Timokratis Karamitros, Gethsimani Papadopoulou, Maria Bousali, Anastasios Mexias, Sotiris Tsiodras, Andreas Mentis

**Author notes:** To whom correspondence should be addressed: Dr. Timokratis Karamitros, Researcher, Unit of Bioinformatics and Applied Genomics, Department of Microbiology, Hellenic Pasteur Institute, 127 Vas. Sophias avenue, 11521, Athens, Greece, m, -874.

## Abstract

In December 2019, an outbreak of atypical pneumonia (Coronavirus disease 2019 - COVID-19) associated with a novel coronavirus (SARS-CoV-2) was reported in Wuhan city, Hubei province, China. The outbreak was traced to a seafood wholesale market and human to human transmission was confirmed. The rapid spread and the death toll of the new epidemic warrants immediate intervention. The intra-host genomic variability of SARS-CoV-2 plays a pivotal role in the development of effective antiviral agents and vaccines, but also in the design of accurate diagnostics.

We analyzed NGS data derived from clinical samples of three Chinese patients infected with SARS-CoV-2, in order to identify small- and large-scale intra-host variations in the viral genome. We identified tens of low- or higher-frequency single nucleotide variations (SNVs) with variable density across the viral genome, affecting 7 out of 10 protein-coding viral genes. The majority of these SNVs corresponded to missense changes. The annotation of the identified SNVs but also of all currently circulating strain variations revealed colocalization of intra-host but also strain specific SNVs with primers and probes currently used in molecular diagnostics assays. Moreover, we de-novo assembled the viral genome, in order to isolate and validate intra-host structural variations and recombination breakpoints. The bioinformatics analysis disclosed genomic rearrangements over poly-A / poly-U regions located in ORF1ab and spike (S) gene, including a potential recombination hot-spot within S gene.

Our results highlight the intra-host genomic diversity and plasticity of SARS-CoV-2, pointing out genomic regions that are prone to alterations. The isolated SNVs and genomic rearrangements, reflect the intra-patient capacity of the polymorphic quasispecies, which may arise rapidly during the outbreak, allowing immunological escape of the virus, offering resistance to anti-viral drugs and affecting the sensitivity of the molecular diagnostics assays.

## INTRODUCTION

Coronaviruses (CoVs), considered to be the largest group of viruses, belong to the *Nidovirales* order, *Coronaviridae* family and *Coronavirinae* subfamily, which is further subdivided into four genera, the alpha- and betacoronaviruses, which infect mammalian species and gamma- and deltacoronaviruses infecting mainly birds [1], [2]. Small mammals (mice, dogs, cats) serve as reservoirs for HCoVs, with significant diversity seen in bats, which are considered to be primordial hosts of HCoVs [3]. On the contrary, peridomestic animals are usually intermediate hosts, who enable long-term establishment of endemicity of the viruses, facilitating mutations and recombination events [1], [4].

Until 2002, minor consideration was given to HCoVs, as they were associated with mild-to-severe disease phenotypes in immunocompetent people [3]–[5]. In 2002, the beginning of severe acute respiratory syndrome (SARS) outbreak took place [6]. In 2005, after the discovery of SARS-CoV-related viruses in horseshoe bats (*Rhinolophus*), palm civets were suggested as intermediate hosts, and bats as primordial hosts of the virus [6], [7]. In 2012, the emerging Middle East respiratory syndrome coronavirus (MERS-CoV) caused an outbreak in Saudi Arabia, which affected both camels and humans (44% mortality).

On December 31^st^ – 2019, a novel Coronavirus (SARS-CoV-2) was first reported from the city of Wuhan, Hubei province in China, causing severe infection of the respiratory tract in humans, after the identification of a group of similar cases of patients with pneumonia of unknown etiology [8] (https://www.who.int/emergencies/diseases/novel-coronavirus-2019). Similarly to SARS, epidemiological links between the majority of 2019-nCoV cases and Huanan South China Seafood Market, a live-animal market, have been reported. A total of 76,775 confirmed cases of “Coronavirus Disease 2019” (COVID-19) were reported up to February 21^st^ 2020, from which 2,247 died and 18,855 recovered. Notably, 75,447 of the confirmed cases were reported in China (https://www.gisaid.org/epiflu-applications/global-cases-betacov/).

The size of the ssRNA genome of SARS-CoV-2 is 29,891 nucleotides, it encodes 9860 amino acids and is characterized by nucleotide identity of ~ 89% with bat SARS-related-CoV SL-ZXC21 and ~ 82% with human SARS-CoVs BJ01 2003 and Tor2 [9]. CoVs are enveloped positive-sense RNA viruses, which are characterized by a very large non-segmented RNA genome (26 to 32kb length), ready to be translated [2], [5]. The genes arrangement on the SARS-CoV-2 genome is: 5′UTR-replicase (ORF1/ab) -Spike (S) -ORF3a -Envelope (E) - Membrane (M) -ORF6 -ORF7a -ORF8 -Nucleocapsid (N) ORF10 −3′UTR [9]. The main difference between SARS-CoV-2 and SARS-CoV is in ORF3b, ORF8 and Spike.

Intra host variability of pathogenic viruses and bacteria represents a significant barrier in the control of infectious diseases. In viral infections, this variation emerges from genomic phenomena taking place during error-prone replication, ending up to multiple circulating quasispecies of low or higher frequency [10], [11]. These variants, in combination with the genetic profile of the host, can potentially influence the natural history of the infection, the viral phenotype, but also the sensitivity of molecular and serological diagnostics assays [12], [13]. Importantly, intra-host genomic variability leads to antigenic variability, which is of higher importance, especially for pathogens that fail to elicit long-lasting immunity in their hosts, and remains a major contributor to the complexity of vaccine design [14], [15]. To date, there are no clinically approved vaccines available for protection of general population from SARS- and MERS-CoV infections as there is no effective vaccine to induce robust cell mediated and humoral immune responses [16], [17].

Here, we explore intra-host genomic variants and low-frequency polymorphic quasispecies in Next Generation Sequencing (NGS) data derived from patients infected by SARS-CoV-2. Our analyses provide insights into the intra-patient pool of viral genomes, identify the frequency levels of rare variants and highlight variable genomic regions and a potential recombination hot-spot within S gene. Intra-host genomic variability is critical for the development of novel drugs and vaccines, which are of urgent necessity, towards the containment of this newly emerging epidemic.

## MATERIALS AND METHODS

In this study we analysed NGS data derived from clinical specimens (oral swabs) from three Chinese patients infected by SARS-CoV-2 (SRA projects PRJNA601736 and PRJNA603194). We aligned the raw read data on reference strain MN975262.1 using bowtie2 [18], after quality check with FastQC (www.bioinformatics.bbsrc.ac.uk/projects/fastqc). The resulting alignments were visualized with the *Integrated Genomics Viewer* (IGV) [19]. After removing PCR duplicates, SNVs were called with a Bonferroni-corrected *P*-value threshold of 0.05 using *samtools* [20] and *LoFreq* [21]. Variants supported by absolute read concordance (>98%) were filtered-out from intra-host variant frequency calculations. We annotated the variations to the reference strain using snpEff [22], SNVs effects were further filtered with snpSift [23] and we estimated the average mutation rate per gene across the viral genome using R scripts. We compared the localization of the intra-host SNVs with all available SNVs observed at population level up to February 18^th^ 2020 (retrieved from www.GISAID.org). We also compared all intra-host and population level SNPs with all primers and probes coordinates to investigate for potential interferences with currently available molecular diagnostic assays [24] (www.who.int/docs/default-source/coronaviruse/peiris-protocol-16-1-20.pdf).

To investigate intra-host genomic rearrangements, we performed *de novo* assembly of the SARS-CoV-2 genomes using Spades [25], and the resulting contigs were analyzed with BLAST [26] and confirmed by remapping of the raw reads. Smaller contigs (<200 bp) were elongated where possible, after pair-wise realignment of the corresponding mapped reads. Basic computations and visualizations we implemented in R programming language R version 3.6.2, using in-house scripts. The secondary structures of the genomic regions surrounding the recombination breakpoints was predicted using RNAfold [27].

## RESULTS

The mapping assembly of the viral genome was almost complete for all samples. The genome coverage and the average read depth across the genome was 99.99% and 133.5x for sample SRR10903401, 99.99% and 522.5x for sample SRR10903402, and 99.94%, and 598.2x for sample SRR10971381, respectively. The alignment statistics for all samples are summarized in **Suppl.Table 1**.

In all cases we isolated the same 5 SNVs with 98-100% read concordance, thus in total divergence with the reference strain (MN975262.1), which were excluded from the downstream analysis. For sample SRR10903401 we isolated 34 lower frequency SNVs in total. Off these, 33 were present with frequencies ranking between 2 and 15%, while only one was present in 40% of the intra-host viral population. The sequencing depth, which is also evaluated during the SNV calling by the LoFreq algorithm, ranked between 39x and 290x at the corresponding SNV positions. The sequencing depth of sample SRR10903402 at the polymorphic positions was substantially higher (103x – 1137x), allowing the isolation of 55 SNVs with frequencies distributed between 0.9% and 14%. The depth over the polymorphic positions of sample SRR10971381 was between 159x – 1872x, allowing the isolation of 10 intra-host SNVs, with frequencies 1.1% - 6.8% (**Figure 1.A, Suppl.Table 2**).

**Figure 1:**
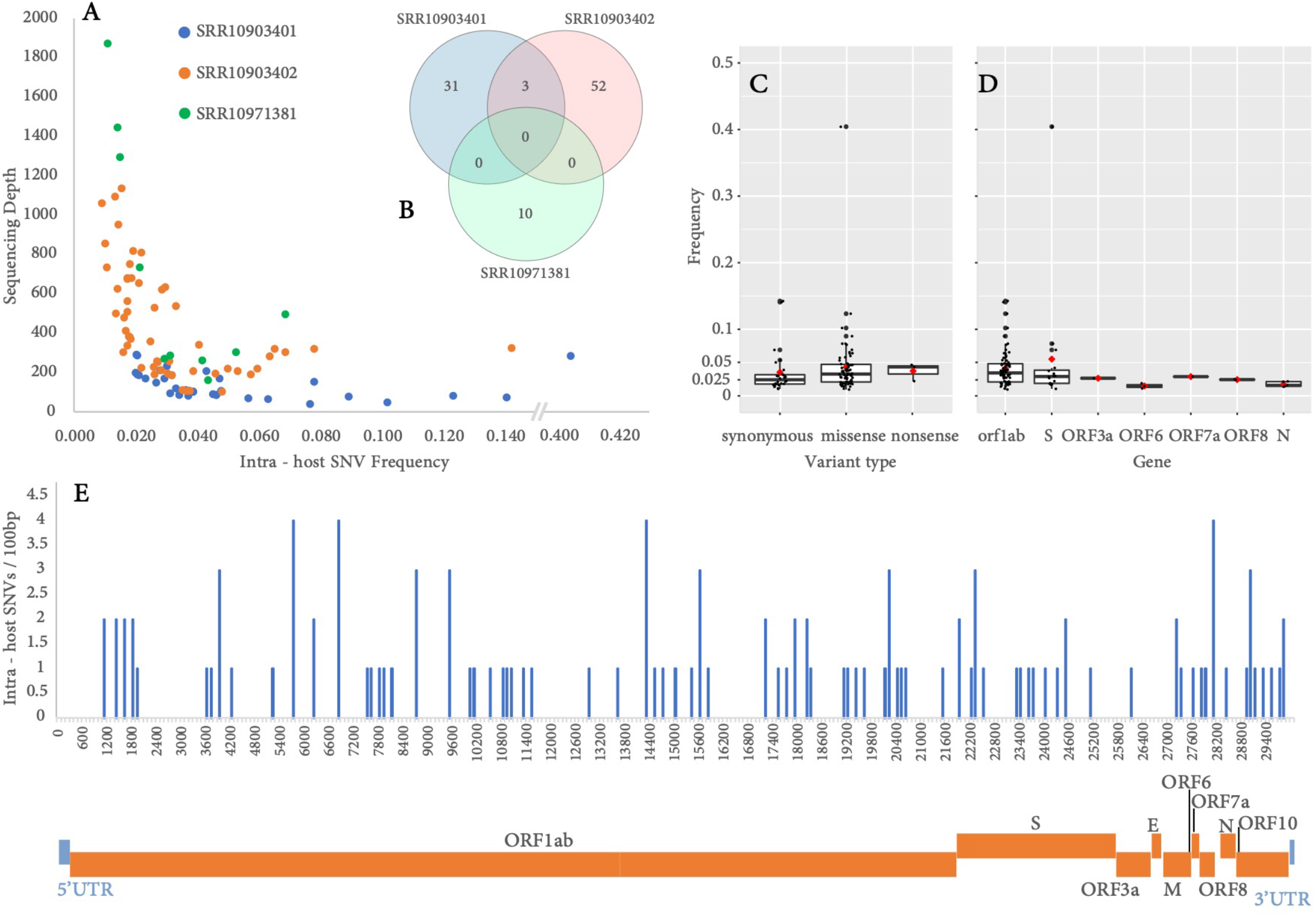
Intra – host SNVs: (A) Intra host SNV frequency vs read depth in the corresponding alignment position (B) SNVs overlaps between the samples analysed (C) Intra-host SNVs frequency vs. variant type – synonymous, missense, monsense (low, moderate, high impact respectively). (D) Intra-host SNVs frequency vs. all seven genes affected (ORF1ab, S, ORF3a, ORF6, ORF7a, ORF8, N). Average values are in red rhombs. (E) Density histogram of intra-host SNVs across the SARS-CoV-2 genome.

Intra-host variants were distributed across 7 out of the 10 protein-coding genes of the viral genome, namely ORF1ab, S, ORF3a, ORF6, ORF7a, ORF8 and N. After normalising for the gene length (variants / kb-gene-length – “v/kbgl”), the higher density was observed in the small ORF6 (16.21 v/kbgl), followed by ORF8 (8.21 v/kbgl), N (4.76 v/kbgl), S (4.18 v/kbgl), ORF1ab (3.47 v/kbgl), ORF7a (2.73 v/kbgl) and ORF3a (1.21 v/kbgl). Interestingly, the majority of the SNPs corresponded to missense changes (leading to amino-acid change) compared to synonymous changes (72 vs. 29 respectively, ratio 2.48:1) (**Table 1**). The average intra-host variant frequency did not differ substantially either between missense and synonymous polymorphisms (**Figure 1.C**), neither between their hosting genes (**Figure 1.D**). We did not detect any small-scale insertions or deletions in the samples (**Suppl. Table 2**).

**Table 1:**
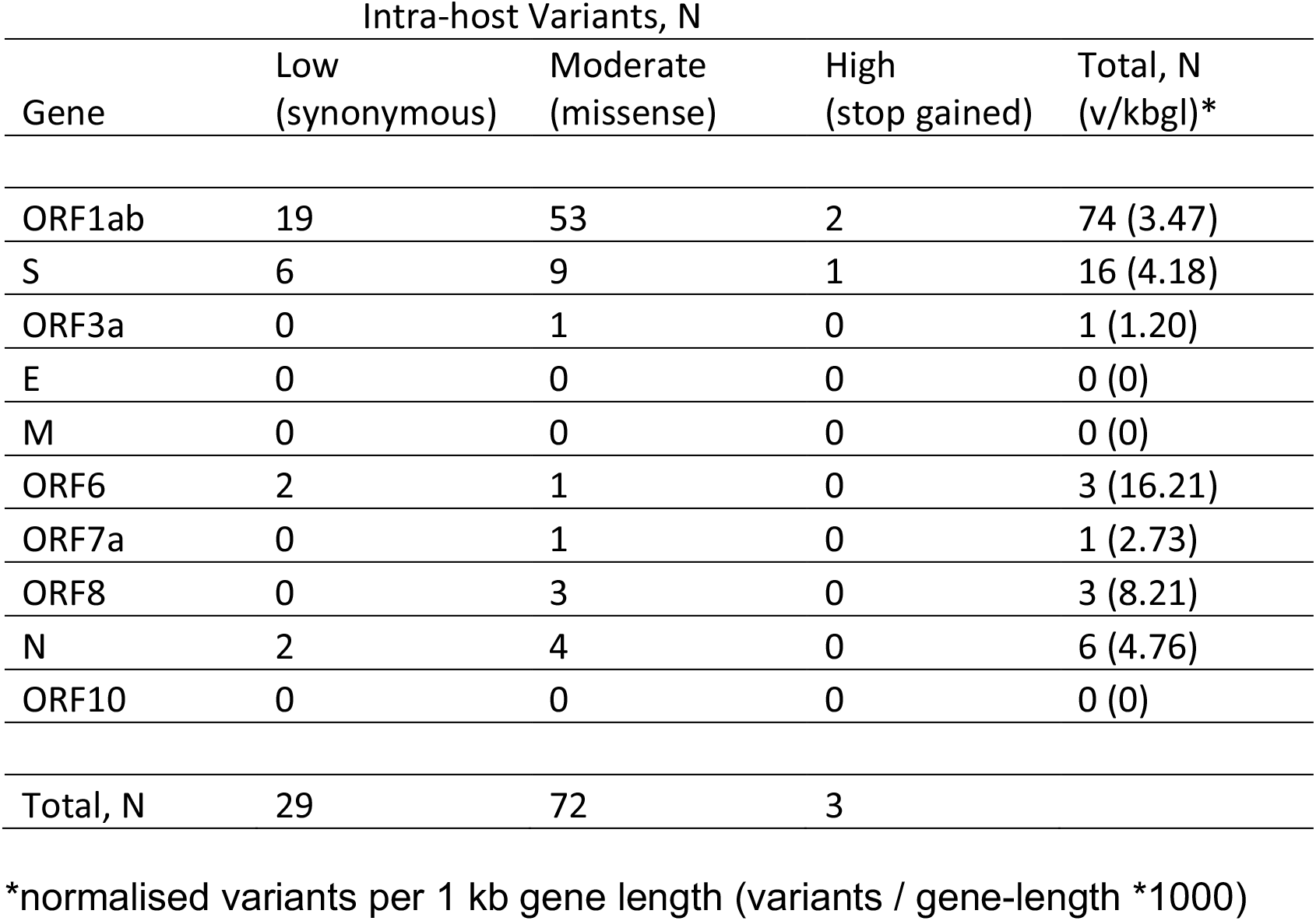
Impact of Intra-host SNVs on viral genes.

The comparison of all SNVs (intra-host and population level) with the genomic targets of the molecular diagnostics assays, revealed colocalizations of three intra-host SNVs and 2 isolate-specific SNVs with primers and probes currently in use. In detail, intra-host SNVs colocalized with the probe of RdRP_SARSr reaction (15,474 T > G), with the reverse primer of HKU-N reaction (28,971 A > G) and with the probe of 2019-nCoV-N2 reaction (29,095 T > C). More importantly, two SNVs belonging to isolates Wuhan/IVD-HB-04/2020 and Chongqing/YC01/2020, colocalized with the forward primer of 2019-nCoV-N1 reaction (28,291 C > T) and the probe of 2019nCoV-N2 reaction (29,200 C > T), respectively (**Figure 2**).

**Figure 2:**
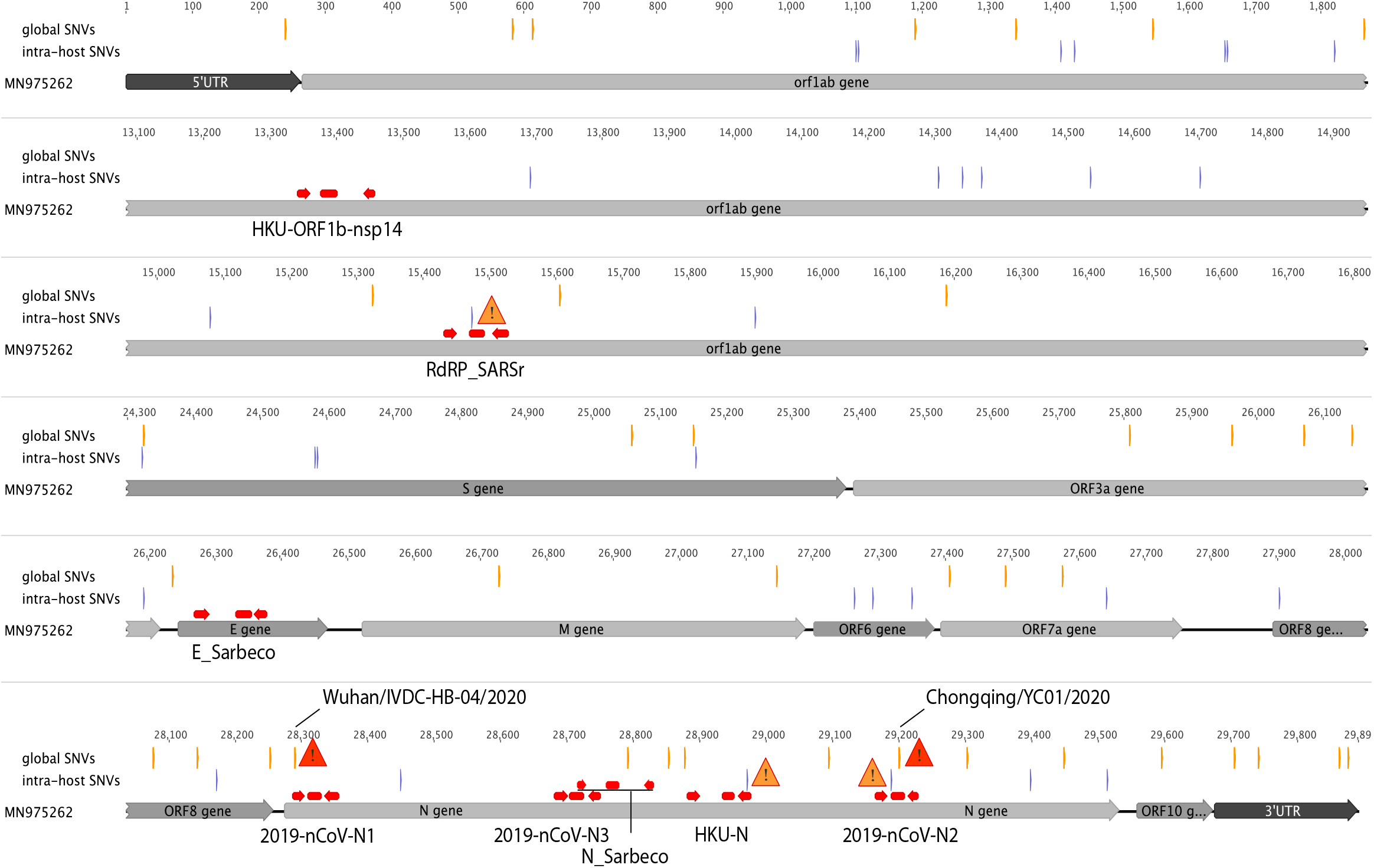
Truncated map of SARS-CoV-2 genome illustrating a subset of intra-host (blue lines) and globally collected, strain-specific SNVs (orange lines) with respect to the genomic targets of molecular diagnostics assays (red arrows – primers, red bars - probes). Three intra-host variants (orange triangles), and two strain specific variants (Wuhan/IVD-HB-04/2020 and Chongqing/YC01/2020 - red triangles), are colocalized with RdRP_SARSr probe (15,474 T > G), HKU-N primer (28,971 A > G) and 2019-nCoV-N2 probe (29,095 T > C).

The *de novo* assembly of the viral genomes revealed intra-host genomic rearrangements. For samples SRR10903401 and SRR10903402, these large-scale structural events were systematically observed over poly-A / poly-U-rich genomic regions, located in ORF1ab and S genes. In all cases, similar or identical strings of nucleotides in close proximity appear to have served as seeds for homologous recombination events. All rearrangements were validated by remapping of the raw reads on the corresponding *de novo* assembled contigs, setting a threshold of at least 5 supporting reads of high mapping quality (>40) in each case. For sample SRR10903401 we isolated three inversions/misassemblies in ORF1ab (**Suppl. Figure 2**) and one inversion/misassembly in S gene (**Figure 3-A**). Notably, we were able to validate the same inversion in S gene for sample SRR10903402 as well (**Figure 3-B**). Apart from 2 inversions in ORF1ab supported by only 2 reads each (not passing the validation threshold), there were no further large-scale intra-host events observed for sample SRR10903402. Similarly, we identified one inversion/misassembly in sample SRR10971381 that was supported by only one read. The alignment coordinates of all rearrangement-supporting contigs with respect to the reference strain are presented in (**Table 2**).

**Figure 3:**
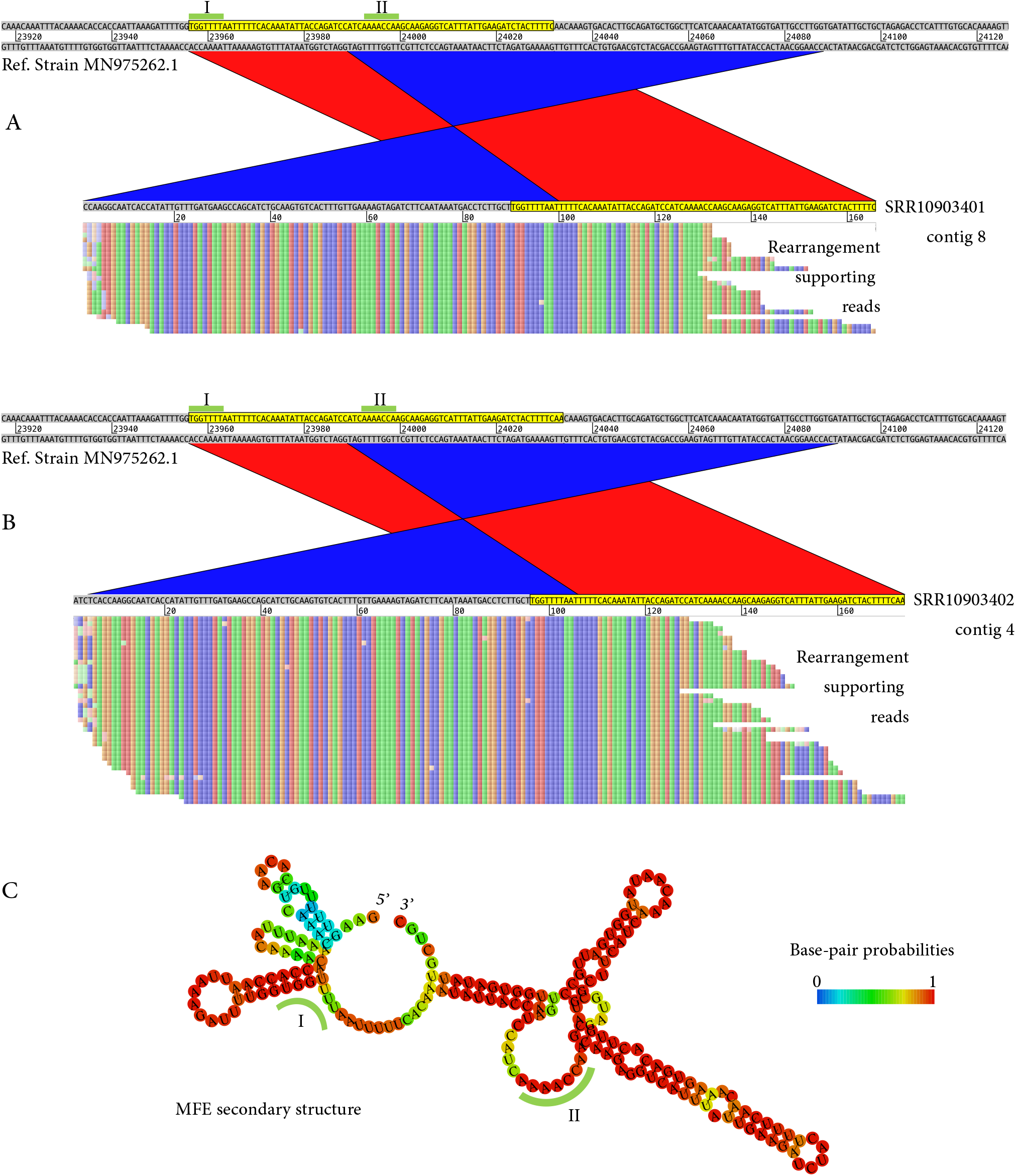
Recombination events in S gene. Samples SRR10903401 (A) and SRR10903402 (B). Alignments of the de novo assembled contigs with respect to the reference genome (MN 975262). Donor – acceptor palindrome sequences are indicated in green bars. Raw, non-duplicated NGS reads, validating the recombination event, are represented below the corresponding contig. (C): Prediction of the secondary structure of the genomic region spanning the rearrangement breakpoint (100 bases upstream and 100 bases downstream). The corresponding donor-acceptor sequences, exposed in internal loops, are indicated in green bars.

**Table 2:** Alignment characteristics of de novo assembled contigs.

| Contig Name | Contig Length | Reference* Coordinates |  | Contig Coordinates |  | Alignment Identity (%) | Alignment Type | Average Read Depth (x) | QC Pass<sup>#</sup> |
| --- | --- | --- | --- | --- | --- | --- | --- | --- | --- |
|  |  | start | end | start | end |  |  |  |  |
| SRR10903401 |  |  |  |  |  |  |  |  |  |
| Contig 1 | 23994 | 75 | 24068 | 23994 | 1 | 99.99 | Correct | 57.01 | + |
| Contig 2 | 5681 | 24246 | 29891 | 1 | 5646 | 99.96 | Correct | 71.40 | + |
| Contig 3 | 331 | 23992 | 24322 | 331 | 1 | 100 | Correct | 164.39 | + |
| Contig 4 | 179 | 24221 | 24399 | 179 | 1 | 100 | Correct | 97.56 | + |
| Contig 5 | 192 | 17816 | 17909 | 94 | 1 | 100 | Inversion | 7.22 | + |
|  |  | 17933 | 18030 | 95 | 192 | 100 | Correct |  |  |
| Contig 6 | 181 | 18052 | 18152 | 101 | 1 | 100 | Relocation, Inconsistency | 8.12 | + |
|  |  | 17766 | 17845 | 102 | 181 | 100 | Misassembly |  |  |
| Contig 7 | 169 | 1707 | 1765 | 62 | 4 | 100 | Inversion | 7.62 | + |
|  |  | 1815 | 1903 | 63 | 151 | 97.75 | Correct |  |  |
| Contig 8 | 165 | 23992 | 24087 | 96 | 1 | 100 | Inversion | 18.04 | + |
|  |  | 23963 | 24031 | 97 | 165 | 100 | misassembly |  |  |
| SRR10903402 |  |  |  |  |  |  |  |  |  |
| Contig 1 | 29842 | 133 | 29891 | 29842 | 84 | 99.98 | Correct | 234.32 | + |
| Contig 2 | 242 | 2075 | 2139 | 178 | 242 | 100 | Partial | 1.09 | - |
| Contig 3 | 242 | 21577 | 21629 | 242 | 190 | 100 | Partial | 1.06 | - |
| Contig 4 | 173 | 23992 | 24090 | 102 | 4 | 100 | Inversion | 39.30 | + |
|  |  | 23963 | 24033 | 103 | 173 | 100 | Misassembly |  |  |
| SRR10971381 |  |  |  |  |  |  |  |  |  |
| Contig 1 | 29902 | 1 | 29891 | 29897 | 7 | 99.98 | Correct | 267.59 | + |
| Contig 2 | 241 | 516 | 559 | 163 | 120 | 100 | Inversion | 1.00 | - |
|  |  | 472 | 501 | 119 | 90 | 100 | Misassembly |  |  |

\* Corresponding to strain MN975262 coordinates
<sup>#</sup> contig supported by at least 5 non duplicated reads of mapping quality >40

## DISCUSSION

The rapid spread and the death toll of the new SARS-CoV-2 epidemic warrants the immediate identification / development of effective antiviral agents and vaccines, but also the design of accurate diagnostics. The intra- and inter-patient variability of the viral genome plays a pivotal role in all the abovementioned efforts, since it affects the compatibility of molecular diagnostics but also impairs the effectiveness of the vaccines and the serological assays by altering the antigenicity of the virus. Intra-host low-frequency variants are also the main source of resistance to anti-viral drugs.

Bioinformatics analysis of NGS data allows the generation of the consensus sequence of a viral genome from the of majority nucleotides at each position but also the identification of non-consensus nucleotides, enabling the exploration of intra-host variability but also its consequences on intra-host viral evolution [28]–[30]. All samples analysed in this study were probably infected by the same viral strain since they shared the same set of consensus SNVs. However, apart from 3 intra-host SNVs that were common between SRR10903401 and SRR10903402, there was no other overlap observed between the low frequency variants of each sample (**Figure 1-B**). This indicates that these variations have been occurred in a rather random fashion and are not subject of selective pressures, which is also supported by the fact that the missense mutations were systematically more, compared to the synonymous mutations. On the other hand, missense substitutions are more common in loci involving pathogen resistance, indicating positive selection [31]. The analysed viral RNA might have been originated from functional/packed virions, but also from unpacked viral genomes, which are unable to replicate and infect other host cells. Even if a viral genome is unable to replicate independently, its abundant presence in the pool of viral quasispecies implies some functionality regarding the intra-host evolution and adaptation. For example, defective viral genomes might affect infection dynamics such as viral persistence but also the natural history of an infection [32]–[34]. At the same time, these variants may arise rapidly during an outbreak and can be used for tracking the transmission chains and the spaciotemporal characteristics of the epidemic [35]–[37]. Studies involving large number of samples and in-vitro experiments on SARS-CoV-2 viral isolates are needed, in order to conclude whether these variations are advantageous or come with a fitness cost for the virus.

SNVs and quasispecies that are observed at low frequency could represent viral variations of low impact on the functionality of the genome. However, their abundance is largely affected by the population size and the epidemic characteristics. For example, a neutral substitution in a region that represents a primer target for a molecular diagnostic assay can drift to fixation rather quickly in a rapidly spreading virus, jeopardizing the sensitivity of the assay [38], [39]. Here, we highlight three intra-host but also two fixed variants that colocalized with primers or probes of real-time PCR diagnostics assays that are currently in use (**Figure 2**). Since the alignment of these oligos with their genomic targets is directly linked to the performance of the corresponding diagnostic assays, the community should pay extra attention in the evaluation of these potentially emerging variations and be alerted, in case redesigning of these oligos is needed.

As it is well documented, recombination events lead to substantial changes in genetic diversity of RNA viruses [40], [41]. In CoVs, discontinuous RNA synthesis is commonly observed, resulting in high frequencies of homologous recombination [42], which can be up to 25% across the entire CoV genome [43]. For pathogenic HCoVs genomic rearrangements are frequently reported during the course of epidemic outbreaks, such as HCoV-OC43 [44], and HCoV-NL63 [45], SARS-CoV [46] [44] and MERS-CoV [47]. We have isolated intra-host genomic rearrangements, located in poly-A and poly-U enriched palindrome regions across the SARS-CoV-2 genome (**Figure 3, Suppl. Figure 1**). We have validated the majority of these events by visual inspection of the alignments. We conclude that these rearrangements do not represent artifacts derived from the NGS library preparation (e.g. PCR crosstalk artifacts), especially since all the supporting reads were not duplicated and, in some cases, differed in polymorphic positions (**Suppl. Figure 2**).

Recombination processes involving S gene particularly, have been reported for SARS- and SARS-like CoV but also for HCoV-OC43. In the case of sister species HCoV-NL63 and HCoV-229E, recombination breakpoints are located near 3’- and 5’-end of the gene [1] [47]. S is a trimeric protein, which is cleaved into two subunits, the globular N-terminal S1 and the C-terminal S2 [48]. The S1 subunit consists of a signal peptide and the NT and receptor binding (RB) domains, with the latter sharing only 40% amino acid identity with other SARS-related CoVs. Our analysis revealed that similarly to other genomic regions, the S1 subunit hosts many low-frequency SNVs, characterized by higher density compared to the rest of the S gene sequence (**Figure 1-E**). The S2 subunit is highly conserved, with 99% identity compared to human SARS-CoV and two bat SARS-like CoVs [9]. The S2 subunit consists of two fusion peptides (FP, IFP), followed by two heptad repeats (HR 1 and 2), the pretransmembrane domain (PTM), the transmembrane and the cytoplasmic domain (TM, CP) [48]. In S gene, the same rearrangement event has taken place in two samples analyzed in this study. This observation highlights a potential recombination hot-spot in S gene. The rearrangement that was common between the two samples of this study is located in nt24,000 of the 2019-nCoV genome, which corresponds to the ~200nt linking region between the fusion peptides FP and IFP (aa 812-813). Examining closely the secondary structure of the RNA genome around the breakpoints, we suggest a model where the palindromes 5’-UGGUUUU-3’ and 5’-AAAACCAA-3’, have served as donor-acceptor sequences during the recombination event, since they are both exposed in the single-stranded internal loops formed in a highly structured RNA pseudoknot (**Figure 3-C**). The RB domain of the S protein has been tested as a potential immunogen as it contains neutralization epitopes which appear to have a role in the induction of neutralizing antibodies [16], [49]. It should be mentioned though that S protein of SARS-CoV is the most divergent in all strains infecting humans [50], [51], as in both C and N-terminal domains variations arise rapidly, allowing immunological escape [52]. Our findings support that apart from these variations, the N-terminal region also hosts a recombination hot-spot, which together with the rest of the observed rearrangements, indicates the genomic instability of SARS-CoV-2 over poly-A and poly-U regions.

## Authors’ contributions

TK: Conceptualization; Data curation; Formal analysis; Methodology; Supervision; Validation; Visualization; Writing - original draft; Writing - review & editing. GP: Data curation; Formal analysis; Writing - original draft; Writing - review & editing. MB: Visualization; Writing - review & editing. AM, ST and AM: Writing - original draft; Writing - review & editing.

## Ethics approval and consent to participate

Not applicable.

## REFERENCES

[1] V. M. Corman, D. Muth, D. Niemeyer, and C. Drosten, “Hosts and Sources of Endemic Human Coronaviruses,” vol. 100, pp. 163–188, 2018, doi: 10.1016/bs.aivir.2018.01.001.

[2] A. R. Fehr and S. Perlman, “HHS Public Access,” pp. 1–23, 2016, doi: 10.1007/978-1-4939-2438-7.

[3] D. Vijaykrishna, G. J. D. Smith, J. X. Zhang, J. S. M. Peiris, H. Chen, and Y. Guan, “Evolutionary Insights into the Ecology of Coronaviruses,” J. Virol., vol. 81, no. 8, pp. 4012–4020, 2007, doi: 10.1128/jvi.02605-06.

[4] C. I. Paules, “Coronavirus Infections - More Than Just the Common Cold,” vol. 2520, pp. 3–4, 2020, doi: 10.1007/82.

[5] F. Li, W. Li, M. Farzan, and S. C. Harrison, “Structural biology: Structure of SARS coronavirus spike receptor-binding domain complexed with receptor,” Science (80-.)., vol. 309, no. 5742, pp. 1864–1868, 2005, doi: 10.1126/science.1116480.

[6] J. Cui, “Origin and evolution of pathogenic coronaviruses,” Nat. Rev. Microbiol., vol. 17, no. March, pp. 181–192, 2019, doi: 10.1038/s41579-018-0118-9.

[7] J. H. Epstein et al., “Bats Are Natural Reservoirs of SARS-Like Coronaviruses,” Science (80-.)., vol. 310, no. 5748, pp. 676–679, 2005, doi: 10.1109/NEMS.2006.334722.

[8] “Novel coronavirus 2019;-nCoV: early estimation of epidemiological parameters and epidemic predictions,” vol. 2020, no. January, 2020.

[9] J. F. Chan et al., “Genomic characterization of the 2019 novel human-pathogenic coronavirus isolated from a patient with atypical pneumonia after visiting Wuhan,” vol. 1751, 2020, doi: 10.1080/22221751.2020.1719902.

[10] B. Li et al., “Rapid Reversion of Sequence Polymorphisms Dominates Early Human Immunodeficiency Virus Type 1 Evolution,” J. Virol., vol. 81, no. 1, pp. 193–201, 2007, doi: 10.1128/jvi.01231-06.

[11] A. Beloukas et al., “Assessment of phylogenetic sensitivity for reconstructing HIV-1 epidemiological relationships,” Virus Res., vol. 166, no. 1–2, pp. 54–60, 2012.

[12] M. Vignuzzi, J. K. Stone, J. J. Arnold, C. E. Cameron, and R. Andino, “Quasispecies diversity determines pathogenesis through cooperative interactions in a viral population,” Nature, vol. 439, no. 7074, pp. 344–348, 2006, doi: 10.1038/nature04388.

[13] T. Karamitros et al., “The interferon receptor-1 promoter polymorphisms affect the outcome of Caucasians with HB eAg-negative chronic HBV infection,” Liver Int., vol. 35, no. 12, pp. 2506–2513, 2015.

[14] R. Malley, K. Trzcinski, A. Srivastava, C. M. Thompson, P. W. Anderson, and M. Lipsitch, “CD4+ T cells mediate antibody-independent acquired immunity to pneumococcal colonization,” Proc. Natl. Acad. Sci. U. S. A., vol. 102, no. 13, pp. 4848–4853, 2005, doi: 10.1073/pnas.0501254102.

[15] M. Lipsitch and J. J. O’Hagan, “Patterns of antigenic diversity and the mechanisms that maintain them,” J. R. Soc. Interface, vol. 4, no. 16, pp. 787–802, 2007, doi: 10.1098/rsif.2007.0229.

[16] C. Y. Yong, H. K. Ong, S. K. Yeap, K. L. Ho, and W. S. Tan, “Recent Advances in the Vaccine Development Against Middle East Respiratory Syndrome-Coronavirus,” Front. Microbiol., vol. 10, no. August, pp. 1–18, 2019, doi: 10.3389/fmicb.2019.01781.

[17] R. J. Nicolas W. Cortes-Penfield, Barbara W. Trautner, “A decade after SARS: Strategies to control emerging coronaviruses,” Physiol. Behav., vol. 176, no. 5, pp. 139–148, 2017, doi: 10.1016/j.physbeh.2017.03.040.

[18] B. Langmead and S. L. Salzberg, “Fast gapped-read alignment with Bowtie 2,” Nat. Methods, vol. 9, no. 4, pp. 357–359, Apr. 2012, doi: 10.1038/nmeth.1923.

[19] J. T. Robinson, H. ThorvaldsdÃóttir, A. M. Wenger, A. Zehir, and J. P. Mesirov, “Variant review with the integrative genomics viewer,” Cancer Research, vol. 77, no. 21. American Association for Cancer Research Inc., pp. e31-e34, 01-Nov-2017, doi: 10.1158/0008-5472.CAN-17-0337.

[20] H. Li et al., “The Sequence Alignment/Map format and SAMtools,” Bioinformatics, vol. 25, no. 16, pp. 2078–2079, Aug. 2009, doi: 10.1093/bioinformatics/btp352.

[21] A. Wilm et al., “LoFreq: A sequence-quality aware, ultra-sensitive variant caller for uncovering cell-population heterogeneity from high-throughput sequencing datasets,” Nucleic Acids Res., vol. 40, no. 22, pp. 11189–11201, Dec. 2012, doi: 10.1093/nar/gks918.

[22] P. Cingolani et al., “A program for annotating and predicting the effects of single nucleotide polymorphisms, SnpEff: SNPs in the genome of Drosophila melanogaster strain w1118; iso-2; iso-3,” Fly (Austin)., vol. 6, no. 2, pp. 80–92, 2012, doi: 10.4161/fly.19695.

[23] P. Cingolani et al., “Using Drosophila melanogaster as a model for genotoxic chemical mutational studies with a new program, SnpSift,” Front. Genet., vol. 3, no. MAR, 2012, doi: 10.3389/fgene.2012.00035.

[24] V. M. Corman et al., “Detection of 2019 novel coronavirus (2019-nCoV) by real-time RT-PCR,” Euro Surveill., vol. 25, no. 3, Jan. 2020, doi: 10.2807/1560-7917.ES.2020.25.3.2000045.

[25] A. Bankevich et al., “SPAdes: A new genome assembly algorithm and its applications to single-cell sequencing,” J. Comput. Biol., vol. 19, no. 5, pp. 455–477, May 2012, doi: 10.1089/cmb.2012.0021.

[26] S. F. Altschul, W. Gish, W. Miller, E. W. Myers, and D. J. Lipman, “Basic local alignment search tool,” J. Mol. Biol., vol. 215, no. 3, pp. 403–410, Oct. 1990, doi: 10.1016/S0022-2836(05)80360-2.

[27] A. R. Gruber, R. Lorenz, S. H. Bernhart, R. Neuböck, and I. L. Hofacker, “The Vienna RNA websuite,” Nucleic Acids Res., vol. 36, no. Web Server issue, p. W70, 2008, doi: 10.1093/nar/gkn188.

[28] M. Cotten et al., “Full-genome deep sequencing and phylogenetic analysis of novel human betacoronavirus,” Emerg. Infect. Dis., vol. 19, no. 5, pp. 736–742, May 2013, doi: 10.3201/eid1905.130057.

[29] B. L. Haagmans, A. C. Andeweg, and A. D. M. E. Osterhaus, “The Application of Genomics to Emerging Zoonotic Viral Diseases,” PLoS Pathog., vol. 5, no. 10, p. e1000557, Oct. 2009, doi: 10.1371/journal.ppat.1000557.

[30] E.-G. Kostaki et al., “Unravelling the history of hepatitis B virus genotypes A and D infection using a full-genome phylogenetic and phylogeographic approach,” Elife, vol. 7, p. e36709, 2018.

[31] D. N. Cooper and N. P. Group., Nature encyclopedia of the human genome. London; New York: Nature Pub. Group, 2003.

[32] J. Aaskov, K. Buzacott, H. M. Thu, K. Lowry, and E. C. Holmes, “Long-term transmission of defective RNA viruses in humans and Aedes mosquitoes,” Science (80-.)., vol. 311, no. 5758, pp. 236–238, Jan. 2006, doi: 10.1126/science.1115030.

[33] P. Farci et al., “The outcome of acute hepatitis C predicted by the evolution of the viral quasispecies,” Science (80-.)., vol. 288, no. 5464, pp. 339–344, Apr. 2000, doi: 10.1126/science.288.5464.339.

[34] E. Karamichali et al., “HCV defective genomes promote persistent infection by modulating the viral life cycle,” Front. Microbiol., vol. 9, p. 2942, 2018.

[35] G. Magiorkinis et al., “HIV-1 epidemic in Russia: an evolutionary epidemiology analysis,” Lancet, vol. 383, p. S71, 2014.

[36] D. Paraskevis et al., “Molecular investigation of HIV-1 cross-group transmissions during an outbreak among people who inject drugs (2011--2014) in Athens, Greece,” Infect. Genet. Evol., vol. 62, pp. 11–16, 2018.

[37] G. Magiorkinis et al., “An innovative study design to assess the community effect of interventions to mitigate HIV epidemics using transmission-chain phylodynamics,” Am. J. Epidemiol., vol. 187, no. 12, pp. 2615–2622, 2018.

[38] J. Y. Noh et al., “Simultaneous detection of severe acute respiratory syndrome, Middle East respiratory syndrome, and related bat coronaviruses by real-time reverse transcription PCR,” Arch. Virol., vol. 162, no. 6, pp. 1617–1623, Jun. 2017, doi: 10.1007/s00705-017-3281-9.

[39] T. Karamitros et al., “A contaminant-free assessment of Endogenous Retroviral RNA in human plasma,” Nat. Sci. Reports, vol. 6, p. 33598, 2016.

[40] E. C. Holmes, The evolution and emergence of RNA viruses. Oxford University Press, 2009.

[41] E.-G. Kostaki et al., “Spatiotemporal characteristics of the HIV-1 CRF02_AG/CRF63_02A1 epidemic in Russia and Central Asia,” AIDS Res. Hum. Retroviruses, vol. 34, no. 5, pp. 415–420, 2018.

[42] M. M. C. Lai, “RNA recombination in animal and plant viruses,” Microbiological Reviews, vol. 56, no. 1. pp. 61–79, 1992, doi: 10.1128/mmbr.56.1.61-79.1992.

[43] R. S. Baric, K. Fu, M. C. Schaad, and S. A. Stohlman, “Establishing a genetic recombination map for murine coronavirus strain A59 complementation groups,” Virology, vol. 177, no. 2, pp. 646–656, Aug. 1990, doi: 10.1016/0042-6822(90)90530-5.

[44] S. K. P. Lau et al., “Molecular Epidemiology of Human Coronavirus OC43 Reveals Evolution of Different Genotypes over Time and Recent Emergence of a Novel Genotype due to Natural Recombination,” J. Virol., vol. 85, no. 21, pp. 11325–11337, Nov. 2011, doi: 10.1128/jvi.05512-11.

[45] K. Pyrc et al., “Mosaic Structure of Human Coronavirus NL63, One Thousand Years of Evolution,” J. Mol. Biol., vol. 364, no. 5, pp. 964–973, Dec. 2006, doi: 10.1016/j.jmb.2006.09.074.

[46] C.-C. Hon et al., “Evidence of the Recombinant Origin of a Bat Severe Acute Respiratory Syndrome (SARS)-Like Coronavirus and Its Implications on the Direct Ancestor of SARS Coronavirus,” J. Virol., vol. 82, no. 4, pp. 1819–1826, Feb. 2008, doi: 10.1128/jvi.01926-07.

[47] J. S. M. Sabir et al., “Co-circulation of three camel coronavirus species and recombination of MERS-CoVs in Saudi Arabia,” Science (80-.)., vol. 351, no. 6268, pp. 81–84, Jan. 2016, doi: 10.1126/science.aac8608.

[48] G. Lu, Q. Wang, and G. F. Gao, “Bat-to-human: Spike features determining ‘host jump’ of coronaviruses SARS-CoV, MERS-CoV, and beyond,” Trends Microbiol., vol. 23, no. 8, pp. 468–478, 2015, doi: 10.1016/j.tim.2015.06.003.

[49] S. Agnihothram et al., “Evaluation of serologic and antigenic relationships between middle eastern respiratory syndrome coronavirus and other coronaviruses to develop vaccine platforms for the rapid response to emerging coronaviruses,” J. Infect. Dis., vol. 209, no. 7, pp. 995–1006, 2014, doi: 10.1093/infdis/jit609.

[50] H. Bisht et al., “Severe acute respiratory syndrome coronavirus spike protein expressed by attenuated vaccinia virus protectively immunizes mice,” Proc. Natl. Acad. Sci. U. S. A., vol. 101, no. 17, pp. 6641–6646, Apr. 2004, doi: 10.1073/pnas.0401939101.

[51] L. Enjuanes, M. L. DeDiego, E. Ãlvarez, D. Deming, T. Sheahan, and R. Baric, “Vaccines to prevent severe acute respiratory syndrome coronavirus-induced disease,” Virus Res., vol. 133, no. 1, pp. 45–62, Apr. 2008, doi: 10.1016/j.virusres.2007.01.021.

[52] L. Jiaming et al., “The recombinant N-terminal domain of spike proteins is a potential vaccine against Middle East respiratory syndrome coronavirus (MERS-CoV) infection,” Vaccine, vol. 35, no. 1, pp. 10–18, Jan. 2017, doi: 10.1016/j.vaccine.2016.11.064.

