## Supplementary Material for "SARS-CoV-2 exhibits intra-host genomic plasticity and low-frequency polymorphic quasispecies"

### Suppl. Table 1: NGS read alignment statistics

| | Sample | | |
|---|---|---|---|
| | SRR10903401 | SRR10903402 | SRR10971381 |
| Paired Reads N (%) | | | |
| Total Number | 476,632 (100) | 676,694 (100) | 28,282,964 (100) |
| Aligned | 13,913 (2.94) | 54,723 (8.18) | 62,288 (0.22) |
| Concordantly Aligned | 11,469 (2.40) | 44,176 (6.52) | 59,261(0.21) |
| Discordantly Aligned | 2444 (0.53) | 10,547 (1.67) | 3027 (0.01) |
| Single Mates N (%) | | | |
| Aligned | 244 (0.03) | 1308 (0.11) | 294(0.001) |
| % Overall Alignment Rate | 2.94 | 8.18 | 0.22 |

### Suppl. Table 2: Annotation of all isolated SNVs on the viral genome (MN975262)

| **SRR10903401** | | | | | | | | | | | |
|---|---|---|---|---|---|---|---|---|---|---|---|
| Position | Ref | Alt | Qual. | Filter Pass | Read Depth | Variant Frequency | Variant Type** | Variant Impact*** | Gene | Nt Modification | AA Modification |
| 1409 | C | T | 106 | + | 104 | 0.048 | MS | M | ORF1ab | c.1144C>T | p.His382Tyr |
| 1821 | G | A | 158 | + | 81 | 0.123 | MS | M | ORF1ab | c.1556G>A | p.Gly519Asp |
| 3695 | C | T | 75 | + | 147 | 0.027 | SV | L | ORF1ab | c.3430C>T | p.Leu1144Leu |
| 3917 | A | G | 67 | + | 86 | 0.047 | MS | M | ORF1ab | c.3652A>G | p.Ser1218Gly |
| 3920 | A | T | 68 | + | 86 | 0.047 | SG | H | ORF1ab | c.3655A>T | p.Lys1219* |
| 3921 | A | T | 67 | + | 88 | 0.045 | MS | M | ORF1ab | c.3656A>T | p.Lys1219Ile |
| 5210 | G | T | 113 | + | 78 | 0.090 | MS | M | ORF1ab | c.4945G>T | p.Ala1649Ser |
| 5702 | C | T | 172 | + | 207 | 0.043 | SG | H | ORF1ab | c.5437C>T | p.Gln1813* |
| 7694 | T | C | 71 | + | 119 | 0.034 | MS | M | ORF1ab | c.7429T>C | p.Phe2477Leu |
| 8782 | T | C | 2433 | + | 67 | *1.000 | SV | L | ORF1ab | c.8517T>C | p.Ser2839Ser |
| 9561 | T | C | 1088 | + | 29 | *1.000 | MS | M | ORF1ab | c.9296T>C | p.Leu3099Ser |
| 10145 | T | C | 79 | + | 106 | 0.038 | MS | M | ORF1ab | c.9880T>C | p.Trp3294Arg |
| 10552 | A | G | 70 | + | 168 | 0.030 | SV | L | ORF1ab | c.10287A>G | p.Glu3429Glu |
| 10817 | G | A | 74 | + | 63 | 0.063 | MS | M | ORF1ab | c.10552G>A | p.Ala3518Thr |
| 10967 | T | C | 60 | + | 95 | 0.032 | MS | M | ORF1ab | c.10702T>C | p.Phe3568Leu |
| 12972 | A | G | 65 | + | 102 | 0.039 | MS | M | ORF1ab | c.12707A>G | p.Asn4236Ser |
| 14537 | T | C | 61 | + | 87 | 0.034 | MS | M | ORF1ab | c.14273T>C | p.Leu4758Pro |
| 14701 | G | A | 63 | + | 39 | 0.077 | MS | M | ORF1ab | c.14437G>A | p.Asp4813Asn |
| 15080 | C | A | 120 | + | 49 | 0.102 | MS | M | ORF1ab | c.14816C>A | p.Ala4939Asp |
| 15607 | C | T | 3146 | + | 86 | *0.988 | SV | L | ORF1ab | c.15343C>T | p.Leu5115Leu |
| 17271 | A | G | 62 | + | 187 | 0.021 | SV | L | ORF1ab | c.17007A>G | p.Lys5669Lys |
| 17300 | A | G | 62 | + | 191 | 0.021 | MS | M | ORF1ab | c.17036A>G | p.Tyr5679Cys |
| 17934 | C | A | 75 | + | 290 | 0.021 | SV | L | ORF1ab | c.17670C>A | p.Thr5890Thr |
| 17936 | G | A | 76 | + | 286 | 0.021 | MS | M | ORF1ab | c.17672G>A | p.Arg5891Lys |
| 18253 | A | T | 142 | + | 168 | 0.048 | MS | M | ORF1ab | c.17989A>T | p.Met5997Leu |
| 18376 | A | G | 62 | + | 196 | 0.020 | MS | M | ORF1ab | c.18112A>G | p.Thr6038Ala |
| 19164 | C | T | 223 | + | 71 | 0.141 | SV | L | ORF1ab | c.18900C>T | p.Asp6300Asp |
| 19229 | T | C | 73 | + | 70 | 0.057 | MS | M | ORF1ab | c.18965T>C | p.Ile6322Thr |
| 19623 | T | G | 73 | + | 102 | 0.039 | MS | M | ORF1ab | c.19359T>G | p.Ser6453Arg |
| 20636 | A | G | 68 | + | 109 | 0.037 | MS | M | ORF1ab | c.20372A>G | p.Glu6791Gly |
| 21510 | A | T | 113 | + | 187 | 0.032 | MS | M | ORF1ab | c.21246A>T | p.Glu7082Asp |
| 21959 | T | C | 62 | + | 80 | 0.038 | MS | M | S | c.397T>C | p.Phe133Leu |

| | | | | | | | | | | | |
|---|---|---|---|---|---|---|---|---|---|---|---|
| 22316 | G | A | 171 | + | 153 | 0.078 | MS | M | S | c.754G>A | p.Gly252Ser |
| 23310 | A | G | 93 | + | 212 | 0.028 | MS | M | S | c.1748A>G | p.Glu583Gly |
| 23725 | T | A | 66 | + | 230 | 0.030 | MS | M | S | c.2163T>A | p.Ser721Arg |
| 24323 | A | C | 3240 | + | 277 | 0.404 | MS | M | S | c.2761A>C | p.Lys921Gln |
| 28144 | C | T | 5965 | + | 160 | *1.000 | MS | M | ORF8 | c.251C>T | p.Ser84Leu |
| 28173 | A | G | 74 | + | 169 | 0.024 | MS | M | ORF8 | c.280A>G | p.Lys94Glu |
| 29095 | T | C | 8458 | + | 231 | *0.996 | SV | L | N | c.822T>C | p.Phe274Phe |
| **SRR10903402** | | | | | | | | | | | |
| 1101 | C | T | 224 | + | 319 | 0.078 | MS | M | ORF1ab | c.836C>T | p.Ser279Phe |
| 1104 | T | A | 189 | + | 320 | 0.066 | MS | M | ORF1ab | c.839T>A | p.Ile280Lys |
| 1429 | T | C | 68 | + | 479 | 0.017 | SV | L | ORF1ab | c.1164T>C | p.His388His |
| 1656 | T | A | 69 | + | 379 | 0.018 | MS | M | ORF1ab | c.1391T>A | p.Val464Asp |
| 1659 | G | T | 67 | + | 383 | 0.018 | MS | M | ORF1ab | c.1394G>T | p.Gly465Val |
| 1821 | G | A | 329 | + | 304 | 0.069 | MS | M | ORF1ab | c.1556G>A | p.Gly519Asp |
| 1927 | T | C | 80 | + | 228 | 0.026 | SV | L | ORF1ab | c.1662T>C | p.Thr554Thr |
| 3761 | G | A | 86 | + | 564 | 0.018 | MS | M | ORF1ab | c.3496G>A | p.Val1166Ile |
| 5710 | A | G | 139 | + | 752 | 0.019 | SV | L | ORF1ab | c.5445A>G | p.Glu1815Glu |
| 5765 | G | A | 89 | + | 680 | 0.018 | MS | M | ORF1ab | c.5500G>A | p.Gly1834Ser |
| 5766 | G | C | 87 | + | 674 | 0.018 | MS | M | ORF1ab | c.5501G>C | p.Gly1834Ala |
| 6254 | G | T | 105 | + | 256 | 0.031 | MS | M | ORF1ab | c.5989G>T | p.Ala1997Ser |
| 6255 | C | T | 90 | + | 256 | 0.027 | MS | M | ORF1ab | c.5990C>T | p.Ala1997Val |
| 6810 | T | C | 65 | + | 112 | 0.036 | MS | M | ORF1ab | c.6545T>C | p.Phe2182Ser |
| 6813 | C | G | 71 | + | 108 | 0.037 | MS | M | ORF1ab | c.6548C>G | p.Thr2183Ser |
| 6816 | G | A | 65 | + | 105 | 0.038 | MS | M | ORF1ab | c.6551G>A | p.Arg2184Lys |
| 6828 | C | G | 84 | + | 103 | 0.049 | MS | M | ORF1ab | c.6563C>G | p.Ser2188Cys |
| 7541 | A | C | 220 | + | 341 | 0.041 | MS | M | ORF1ab | c.7276A>C | p.Ile2426Leu |
| 7854 | A | G | 72 | + | 624 | 0.014 | MS | M | ORF1ab | c.7589A>G | p.Asn2530Ser |
| 7970 | A | T | 89 | + | 1058 | 0.009 | MS | M | ORF1ab | c.7705A>T | p.Ile2569Leu |
| 8196 | C | T | 77 | + | 499 | 0.014 | MS | M | ORF1ab | c.7931C>T | p.Ser2644Leu |
| 8782 | T | C | 6957 | + | 201 | *0.980 | SV | L | ORF1ab | c.8517T>C | p.Ser2839Ser |
| 9561 | T | C | 3842 | + | 105 | *1.000 | MS | M | ORF1ab | c.9296T>C | p.Leu3099Ser |
| 10080 | C | T | 71 | + | 410 | 0.017 | MS | M | ORF1ab | c.9815C>T | p.Pro3272Leu |
| 11367 | A | T | 199 | + | 190 | 0.058 | MS | M | ORF1ab | c.11102A>T | p.Tyr3701Phe |
| 11563 | C | T | 1017 | + | 323 | 0.142 | SV | L | ORF1ab | c.11298C>T | p.Cys3766Cys |
| 13693 | A | T | 276 | + | 281 | 0.064 | MS | M | ORF1ab | c.13429A>T | p.Thr4477Ser |
| 14307 | T | C | 142 | + | 219 | 0.050 | SV | L | ORF1ab | c.14043T>C | p.Tyr4681Tyr |
| 14308 | T | C | 91 | + | 217 | 0.060 | MS | M | ORF1ab | c.14044T>C | p.Trp4682Arg |

| | | | | | | | | | | | |
|---|---|---|---|---|---|---|---|---|---|---|---|
| 14344 | T | C | 79 | + | 187 | 0.032 | SV | L | ORF1ab | c.14080T>C | p.Leu4694Leu |
| 14373 | A | G | 67 | + | 225 | 0.022 | SV | L | ORF1ab | c.14109A>G | p.Ala4703Ala |
| 15474 | T | G | 80 | + | 205 | 0.054 | SV | L | ORF1ab | c.15210T>G | p.Gly5070Gly |
| 15607 | C | T | 8759 | + | 238 | *0.992 | SV | L | ORF1ab | c.15343C>T | p.Leu5115Leu |
| 15900 | T | C | 77 | + | 210 | 0.029 | SV | L | ORF1ab | c.15636T>C | p.Val5212Val |
| 17543 | T | A | 76 | + | 732 | 0.011 | MS | M | ORF1ab | c.17279T>A | p.Met5760Lys |
| 17737 | A | G | 168 | + | 819 | 0.020 | MS | M | ORF1ab | c.17473A>G | p.Thr5825Ala |
| 18253 | A | T | 189 | + | 634 | 0.030 | MS | M | ORF1ab | c.17989A>T | p.Met5997Leu |
| 19452 | T | A | 84 | + | 383 | 0.018 | SV | L | ORF1ab | c.19188T>A | p.Ala6396Ala |
| 20236 | A | C | 67 | + | 194 | 0.031 | SV | L | ORF1ab | c.19972A>C | p.Arg6658Arg |
| 20236 | A | G | 122 | + | 194 | 0.046 | MS | M | ORF1ab | c.19972A>G | p.Arg6658Gly |
| 20238 | G | A | 95 | + | 205 | 0.039 | SV | L | ORF1ab | c.19974G>A | p.Arg6658Arg |
| 20412 | A | C | 66 | + | 254 | 0.028 | MS | M | ORF1ab | c.20148A>C | p.Glu6716Asp |
| 21904 | C | T | 65 | + | 304 | 0.016 | SV | L | S | c.342C>T | p.Thr114Thr |
| 22270 | T | C | 124 | + | 509 | 0.018 | SV | L | S | c.708T>C | p.Thr236Thr |
| 22316 | G | A | 218 | + | 539 | 0.033 | MS | M | S | c.754G>A | p.Gly252Ser |
| 22326 | C | T | 157 | + | 528 | 0.027 | MS | M | S | c.764C>T | p.Ser255Phe |
| 23687 | A | G | 96 | + | 679 | 0.019 | MS | M | S | c.2125A>G | p.Asn709Asp |
| 24082 | C | A | 70 | + | 810 | 0.022 | SG | H | S | c.2520C>A | p.Cys840* |
| 24583 | T | C | 74 | + | 856 | 0.011 | SV | L | S | c.3021T>C | p.Tyr1007Tyr |
| 25156 | C | T | 79 | + | 336 | 0.018 | SV | L | S | c.3594C>T | p.Ile1198Ile |
| 26194 | A | T | 72 | + | 189 | 0.026 | MS | M | ORF3a | c.802A>T | p.Thr268Ser |
| 27292 | T | C | 80 | + | 370 | 0.019 | MS | M | ORF6 | c.91T>C | p.Tyr31His |
| 27644 | C | G | 205 | + | 622 | 0.029 | MS | M | ORF7a | c.251C>G | p.Pro84Arg |
| 27904 | T | A | 70 | + | 357 | 0.025 | MS | M | ORF8 | c.11T>A | p.Leu4His |
| 28144 | C | T | 19121 | + | 522 | *1.000 | MS | M | ORF8 | c.251C>T | p.Ser84Leu |
| 28971 | A | G | 99 | + | 1137 | 0.016 | MS | M | N | c.698A>G | p.Lys233Arg |
| 29095 | T | C | 34590 | + | 955 | *0.996 | SV | L | N | c.822T>C | p.Phe274Phe |
| 29188 | A | G | 76 | + | 1093 | 0.014 | SV | L | N | c.915A>G | p.Ala305Ala |
| 29398 | G | C | 109 | + | 950 | 0.015 | MS | M | N | c.1125G>C | p.Lys375Asn |
| 29514 | C | G | 91 | + | 654 | 0.021 | MS | M | N | c.1241C>G | p.Ala414Gly |
| **SRR10971381** | | | | | | | | | | | |
| 4281 | T | C | 181 | + | 302 | 0.052 | MS | M | ORF1ab | c.4016T>C | p.Val1339Ala |
| 27264 | T | C | 76 | + | 1872 | 0.018 | SV | L | ORF6 | c.63T>C | p.Thr21Thr |
| 27351 | T | C | 86 | + | 1446 | 0.014 | SV | L | ORF6 | c.150T>C | p.Ser50Ser |
| 20104 | C | T | 92 | + | 1294 | 0.015 | MS | M | ORF1ab | c.19840C>T | p.Leu6614Phe |
| 28450 | T | A | 70 | + | 732 | 0.021 | MS | M | N | c.177T>A | p.His59Gln |

| | | | | | | | | | | | |
|---|---|---|---|---|---|---|---|---|---|---|---|
| 24586 | G | C | 81 | + | 268 | 0.029 | SV | L | S | c.3024G>C | p.Val1008Val |
| 20542 | T | C | 73 | + | 284 | 0.031 | MS | M | ORF1ab | c.20278T>C | p.Ser6760Pro |
| 22592 | G | A | 88 | + | 262 | 0.041 | MS | M | S | c.1030G>A | p.Ala344Thr |
| 11049 | T | C | 71 | + | 159 | 0.044 | MS | M | ORF1ab | c.10784T>C | p.Val3595Ala |
| 23434 | T | C | 341 | + | 493 | 0.068 | SV | L | S | c.1872T>C | p.Ile624Ile |

*Variants with >98% frequency were excluded from downstream analysis of low-frequency variants.

** Variant Type: MS=missense, SV=synonymous variant, SG= stop gained

***Variant Impact: L=low, M=moderate, H=high

### Suppl. Figure 1

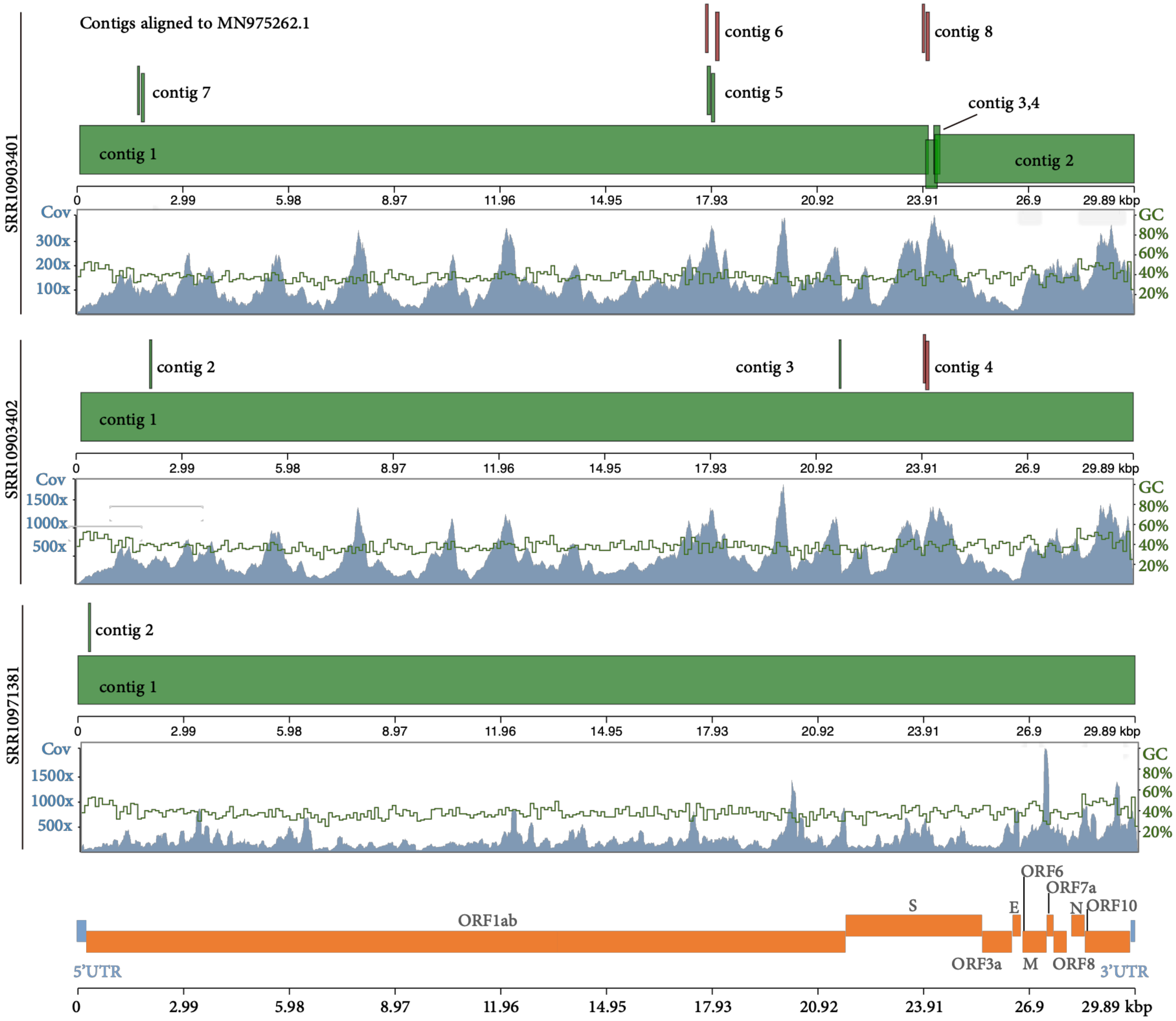


**Suppl. Figure 1:** Alignment of the de novo assembled contigs on the genomic map (bottom). Concordantly aligned contigs (correct or gapped) are in green, while discordantly aligned are in red. Read depth plot (coverage) across the genome (blue) and relative % GC content (green) is presented for each sample.

### Suppl.Figure 2

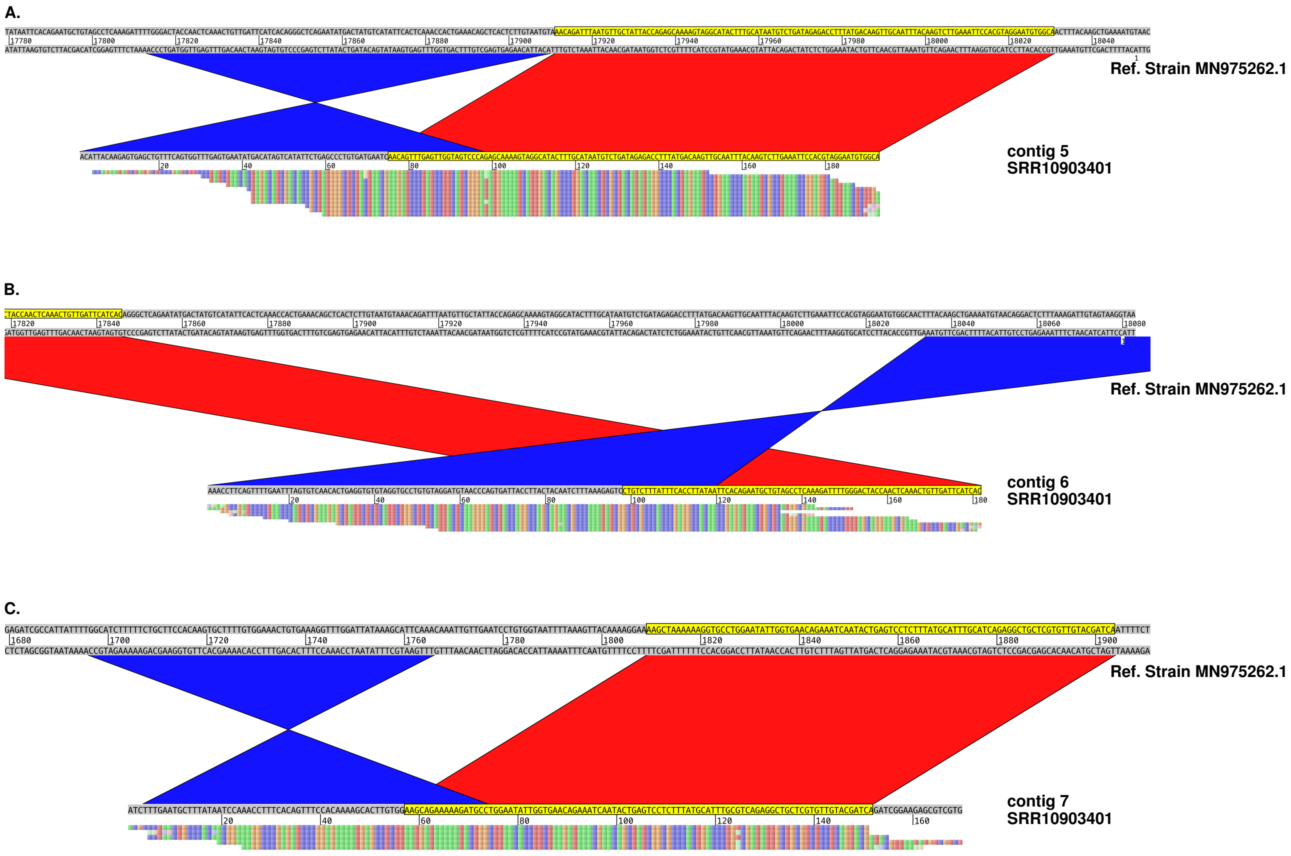


**Suppl.Figure 2:** Recombination events in ORF1ab in sample SRR10903401. Alignments of the de novo assembled contigs with respect to the reference genome (MN 975262). Raw, non-duplicated NGS reads, validating the recombination event, are represented below the corresponding contig.
